# Blue sky’s the limit? Somatic embryogenesis as a means of propagating recalcitrant blue spruce (*Picea pungens*) cultivar Hoopsii

**DOI:** 10.1101/700518

**Authors:** Jordan Demone, Jingqin Mao, Shen Wan, Maryam Nourimand, Äsbjörn Erik Hansen, Barry Flinn, Olivia Facchin, Alar Prost, Illimar Altosaar

**Author notes:** These authors contributed equally to this work (Jordan Demone, Jingqin Mao). **Corresponding author:** Illimar Altosaar.

## Abstract

The ‘triple-blue’ cultivar of blue spruce (*Picea pungens* Hoopsii) is notably recalcitrant towards the realm of traditional vegetative propagation methods. Its ability to naturally proliferate is limited by ovule and embryo abortion during the growing season, leading to low viable seed yield. In this study, we established a protocol using somatic embryogenesis (SE) as a means of propagating this popular ornamental cultivar. We collected cones from Hoopsii trees at seven different timepoints throughout the growing season (mid-June to late July in Ottawa (Plant Hardiness Zone 5A)). Female megagametophytes were harvested following each collection and immature zygotic embryos were plated onto induction media. Early somatic embryos began developing from the embryonic tissue (ET) three to five weeks following induction. The highest ET initiation frequency occurred from embryos collected June 20–July 10, suggesting that developmental stage of the embryo was a significant factor in SE induction. The conversion of mature somatic embryos into plantlets (emblings) was completed in eight–ten weeks at a rate of 92.8%. In this study, we demonstrate that *in vitro* somatic embryogenesis using our optimized protocol is a fast and prolific method for the mass propagation of Hoopsii blue spruce. This is the first report on the production of somatic Hoopsii emblings.

## Introduction

The ‘triple-blue’ spruce (*Picea pungens* Hoopsii) is a popular ornamental conifer. Its attractive silver-blue foliage, compact growth, and pyramidal shape are frequently striking features of the landscape. Like many cultivars of blue spruce, Hoopsii is marketed as an attractive Christmas tree and has become popular across both North American and Europe. Hoopsii is also unique among blue spruces, in that it maintains its silvery coloration for several years longer. However, the high rates of ovule and embryo abortion in Hoopsii results in poor seed yield, and trees that do grow from a viable seed may not grow true. Thus, Hoopsii is an extremely rare tree, and many people have likely never seen one in their lifetime. Despite their high economic value, research on these trees is limited due to its scarcity (Fladung et al. 2012). In addition, the combined threat of climate change (Schlyter et al. 2006) and pests that target spruces (Marini et al. 2017), presses the issue of developing a simple and straightforward means of propagating Hoopsii. To date there have been few investigations into its seed morphology and potential means of highly effective propagation.

Grafting has been the only method for the proliferation of many recalcitrant conifers (Larson 2006; Nanda and Melnyk 2018). Difficulty in repurposing conventional systems for Hoopsii has been a barrier for further economic development. There are two issues that arise in tissue propagation of Hoopsii trees: (1) rooting of Hoopsii cuttings is only possible when stock trees are young (Greenwood 1987; 1995), and Hoopsii trees have already been propagated asexually to the point where cuttings are no longer active; and (2) Hoopsii stem cuttings do not respond to any treatments on propagation media. Thus, plant regeneration via *in vitro* somatic embryogenesis (SE) may be the key to successfully propagating this popular cultivar. SE is a low-cost method for producing genetically uniform plants, particularly for recalcitrant species (Stasolla and Yeung 2003). Conifers are historically recalcitrant towards *in vitro* embryo induction, but there has been recent progress in the use of SE for the initiation of callus or embryonic tissue (ET) from conifers. Since the first report of using somatic embryogenesis to propagate Norway spruce (*Picea abies*) (Hakman and Arnold 1985), several *Picea* species have been successfully propagated via SE, including black spruce (*Picea mariana*) (Adams et al. 1994), white spruce (*Picea glauca*) (Webb et al. 1989), interior spruce (*Picea glauca* x *engelmannii*) (Webster et al. 1990), and Korean spruce (*Picea koraiensis* Nakai) (Li et al. 2008). While successful SE for *P. pungens* has been reported previously for other genotypes (Afele et al. 1992), SE has yet to be reported for the valuable Hoopsii ‘triple-blue’ genotype. The present study evaluates the potential of Hoopsii regeneration via *in vitro* SE and optimizes the factors to develop a feasible and productive protocol for commercial mass propagation of Hoopsii blue spruce.

## Materials and methods

### Plant materials

To collect female megagametophytes and immature zygotic embryos: female *Picea pungens* Hoopsii cones (strobili) were collected from three open-pollinated plots in Ottawa from 2002–2004 (Plant Hardiness Zone 5A (McKenney 2001). Two trees (aged 14 and 25 years old) on residential properties were selected as stock trees. Grove-planted trees (25 years old) standing at the Rideau Valley Conservation Authority in Manotick, Ottawa (45°14’51.3”N 75°42’23.4”W), were also used as stock trees. The cone collection schedule was based on an established conifer embryo development timeline (Klimaszewska et al. 2001). Cones were harvested weekly, biweekly, and monthly from early June to late July (June 12/16, June 18/20/21, June 24/30, July 3/5/6, July 10/14/17, July 17/19, and July 26). The same experimental design was repeated for three years (2002–2004). To obtain immature zygotic embryos: cones were soaked in 0.2% KMnO_4_ for five minutes, rinsed in unsterilized tap water, and surface-sterilized with 70% ethanol for 30 seconds and 15% household bleach for 15 minutes. To obtain embryos from cones collected after June 20: immature zygotic embryos were excised from ovuliferous scales using a sterile scalpel. For cones collected earlier than June 20: translucent female megagametophytes (embryo sacs) containing immature zygotic embryos were collected and plated whole due to the difficulty in extracting intact embryos of this size (<1 mm) from the megagametophyte. Extracted embryos and megagametophytes were placed onto induction media for three to five weeks. Embryonic callus was divided into two groups: Group 1 was transferred to maturation media for somatic embryo development culture. Group 2 was maintained on the same media for continued proliferation of embryonic callus.

### Medium formulations and culture conditions

The basal medium for callus induction contained the following: 0.5X MS (Murashige and Skoog 1962), 1 g/L casein hydrolysate, 500 mg/L filter-sterilized L-glutamine (assists the embryo with nitrogen metabolism (Carlsson et al. 2019)), 20 g/L sucrose, and 4 g/L Phytagel, pH 5.8. Medium was supplemented with 1.0 mg/L 6-benzylaminopurine (BAP) and 4.0 mg/L 2,4-dichlorophenoxyacetic acid (2,4-D), the standard ratio of plant growth regulators in conifer callus induction medium (Carneros et al. 2009; Lara-Chavez et al. 2011; Montalbán et al. 2012). All explants plated onto these media were incubated in darkness at 25±2°C and subcultured onto fresh initiation media every two weeks. For each collection date, three to five Petri dishes (60 x 15 mm) were used, each containing four to six zygotic embryos. The initiation results were monitored using a stereomicroscope during culturing. Initiation frequencies were reported after eight weeks. Initiation frequency was calculated using the following equation:

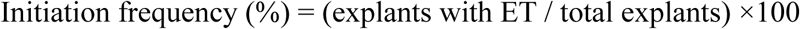

### Maturation and germination of somatic embryos

Maturation and germination of somatic embryos followed the protocol of the previous study on Eastern white pine (*Pinus strobus*) with a few modifications (Klimaszewska et al. 2001). To make 0.5X MS maturation media: filter-sterilized abscisic acid (ABA) was added at 15 mg/L; neither BAP nor 2,4-D were added. Medium osmoticum was also increased by increasing sucrose to 60 g/L and Phytagel to 6 g/L. ET that developed on the surfaces of the callus were excised when embryonal heads and suspensors appeared. Each gram of fresh ET was suspended in 10 mL liquid induction medium in a 15 mL centrifuge tube. The tube was vigorously agitated to form a fine suspension. Suspension liquid was withdrawn with a wide-mouth pipette and poured over a sterile filter paper disk into a Büchner funnel. Liquid was removed by vacuum pulse and the paper disk was transferred onto the same induction medium and subcultured for one week. Maturation culture was performed under low light intensity (5 μM m-2 s-1), with a 16h photoperiod at 25±1°C. The following developmental stages were observed in embryos plated onto maturation media: globular-shaped, torpedo-shaped, cotyledonary (visible cotyledons), and mature somatic embryos (fully developed cotyledons). After six to eight weeks on maturation media, fully mature somatic embryos were transferred onto germination medium (0.5X MS with 20% sucrose and 6 g/L Phytagel). Embryos were placed onto the medium in rows with approximately 30 embryos per plate. Plates were placed vertically in a growth chamber under light conditions of 10 μM m-2 s-1 for the first week, 20 μM m-2 s-1 for the second week, and 30 μM m-2 s-1 until germination occurred. After 30 days of growth, emblings (plantlets) were transferred onto fresh germination medium (15 emblings per plate). Emblings that developed roots were ‘hardened’ two weeks prior to soil transplantation in the greenhouse by removing the Parafilm tape from the Petri dishes. Soil medium was composed of a 3:1:1 ratio of peat, perlite, and vermiculite, respectively.

## Results

### Initiation of embryonic tissue

Initiation responses of *Picea pungens* Hoopsii immature zygotic embryos collected at different developmental stages were investigated in this study. Cones (strobili) were collected at seven timepoints throughout the growing season (June 12/16 to July 26) (Table 1) from adult Hoopsii trees found at multiple locations in Ottawa (Fig. 1a). The experiment was performed three times (2002 growing season to the 2004 growing season). For the first timepoint, cones were collected mid-June. These cones contained translucent megagametophytes (0.5–1.0 mm) and only produced non-ET callus on induction media. Cones collected during the second timepoint (June 18/20/21) contained translucent but larger megagametophytes (1.0–1.5 mm). The embryos harvested from these megagametophytes responded to induction and produced ET callus (Table 1). For the third collection timepoint (June 24/30): megagametophytes were larger (2 mm) and opaque, but the embryos produced ET callus. For the fourth collection timepoint (July 3/5/6): the megagametophytes were opaque and 2–3 mm in size but the embryos still responded to the induction medium by producing ET calls. For the fifth collection timepoint (July 10/14/17): there were virtually no viable embryos remaining. Cones collected on the sixth (July 17/19) and seventh timepoints (July 26) were devoid of viable embryos. The sixth and seventh timepoints were not collected in 2003. There were two types of cell mass produced from immature zygotic embryos when induced: (1) ET cell mass (Fig. 1b) and (2) non-ET cell mass that failed to regenerate into somatic embryos. After three to five weeks on induction culture media, translucent and filamentous tissue emerged from the radicle end of the megagametophyte. Embryo production was characterized by the appearance of embryonal ‘heads’ (composed of meristematic cells (Salaj et al. 2019)) connected by vacuolated ‘suspensors’ emerging from the polyembryonic cell mass (Fig. 1c). When harvested and plated onto maturation media, these tissues developed into mature somatic embryos (Fig. 1d). Cotyledons developed after four additional weeks of growth on maturation media (Fig. 1e). The highest ET initiation frequency attained was 7.67% from precotyledonary immature zygotic embryos collected between roughly June 20 and July 10.

**Table 1.** Embryonic tissue (ET) initiation from immature zygotic embryos from different collection dates

| Timepoint | Collection dates | Explant source characteristics | Initiation results |  |
| --- | --- | --- | --- | --- |
|  |  |  | ET | Non-ET |
| 1 | 6/12/2002<br>6/16/2003<br>6/12/2004 | Translucent megagametophytes: 0.5–1.0 mm | no | yes |
| 2 | 6/20/2002<br>6/21/2003<br>6/18/2004 | Translucent megagametophytes: 1.0–1.5 mm | yes | yes |
| 3 | 6/30/2003<br>6/24/2004 | Opaque megagametophytes: 2 mm | yes | yes |
| 4 | 7/03/2002<br>7/06/2003<br>7/05/2004 | Opaque megagametophytes: 2–3 mm | yes | yes |
| 5 | 7/10/2002<br>7/14/2003<br>7/17/2004 | Hard opaque megagametophytes | yes | yes |
| 6 | 7/19/2002<br>7/17/2004 | No viable embryos | no | no |
| 7 | 7/26/2002<br>7/26/2004 | No viable embryos | no | no |

**Fig. 1.**
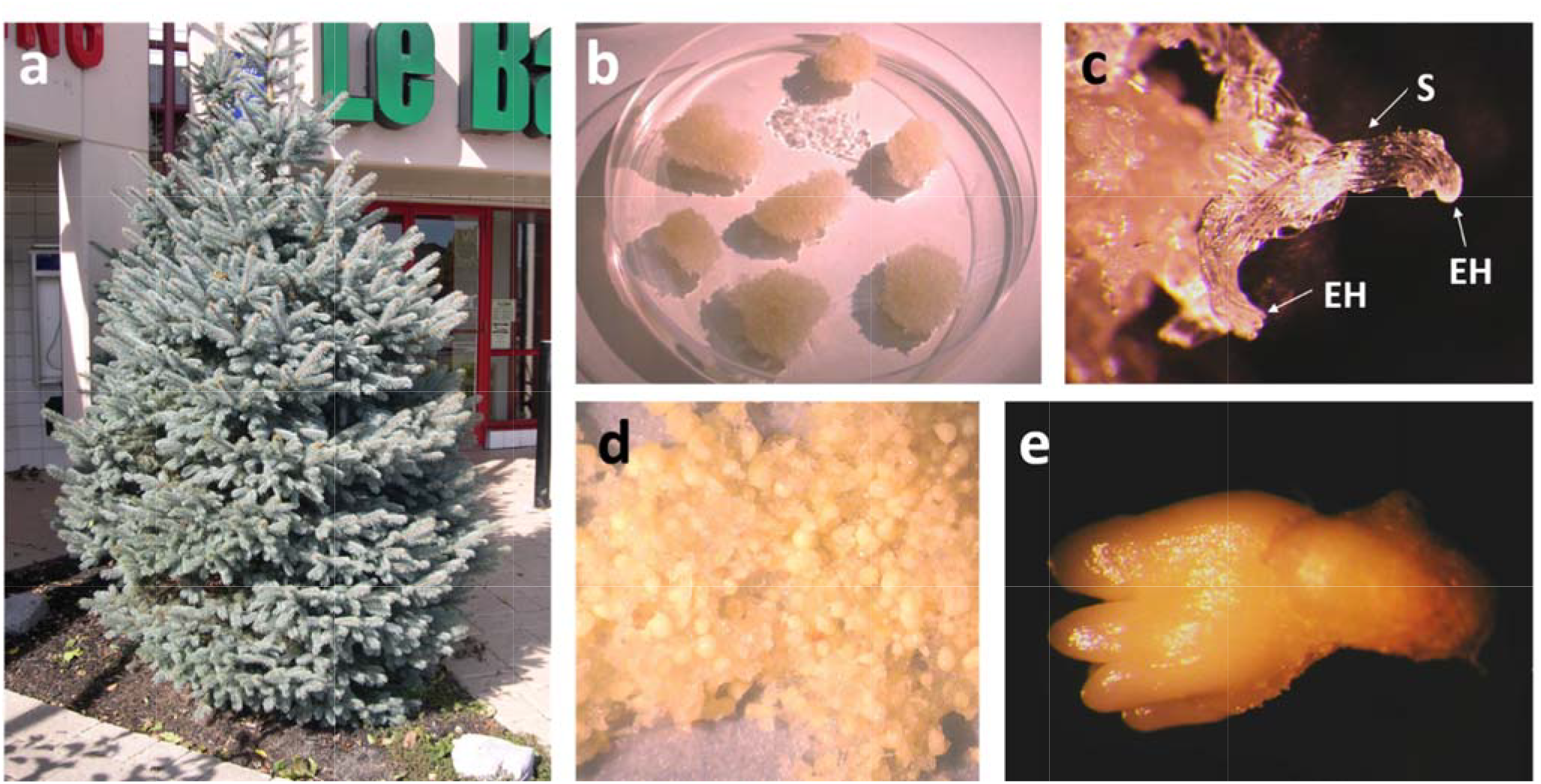
Somatic embryogenesis (SE) as a means of mass propagating *Picea pungens* Hoopsii **a** 10-year-old Hoopsii tree **b** Embryonic tissue (ET) generated by plating immature zygotic embryos onto induction media **c** Initiation of SE tissue (arrow) Suspensors (S) and embryonal heads (EH) are indicated by arrows **d** Maturing somatic embryos after four weeks on maturation culture **e** Cotyledonary embryo

### Maturation culture of somatic embryos

Four distinct stages of embryo development were observed during Hoopsii somatic embryogenesis (Fig. 2): globular-shaped stage, torpedo-shaped stage, cotyledonary (visible cotyledons) and mature somatic embryos (fully developed cotyledons, able to germinate). After two weeks on maturation medium, distinct globular somatic embryos became visible on the surface of the callus. After a further three to four weeks on maturation medium, cotyledonary primordia developed. Early or late cotyledonary stages of embryos were observed around four to five weeks of development. The embryos required another three to four weeks on maturation media to fully mature. By eight to ten weeks, mature cotyledonary embryos had formed (Fig. 3a). Fully developed, healthy somatic embryos accounted for 78.2% of the initial number of immature zygotic embryos.

**Fig. 2.**
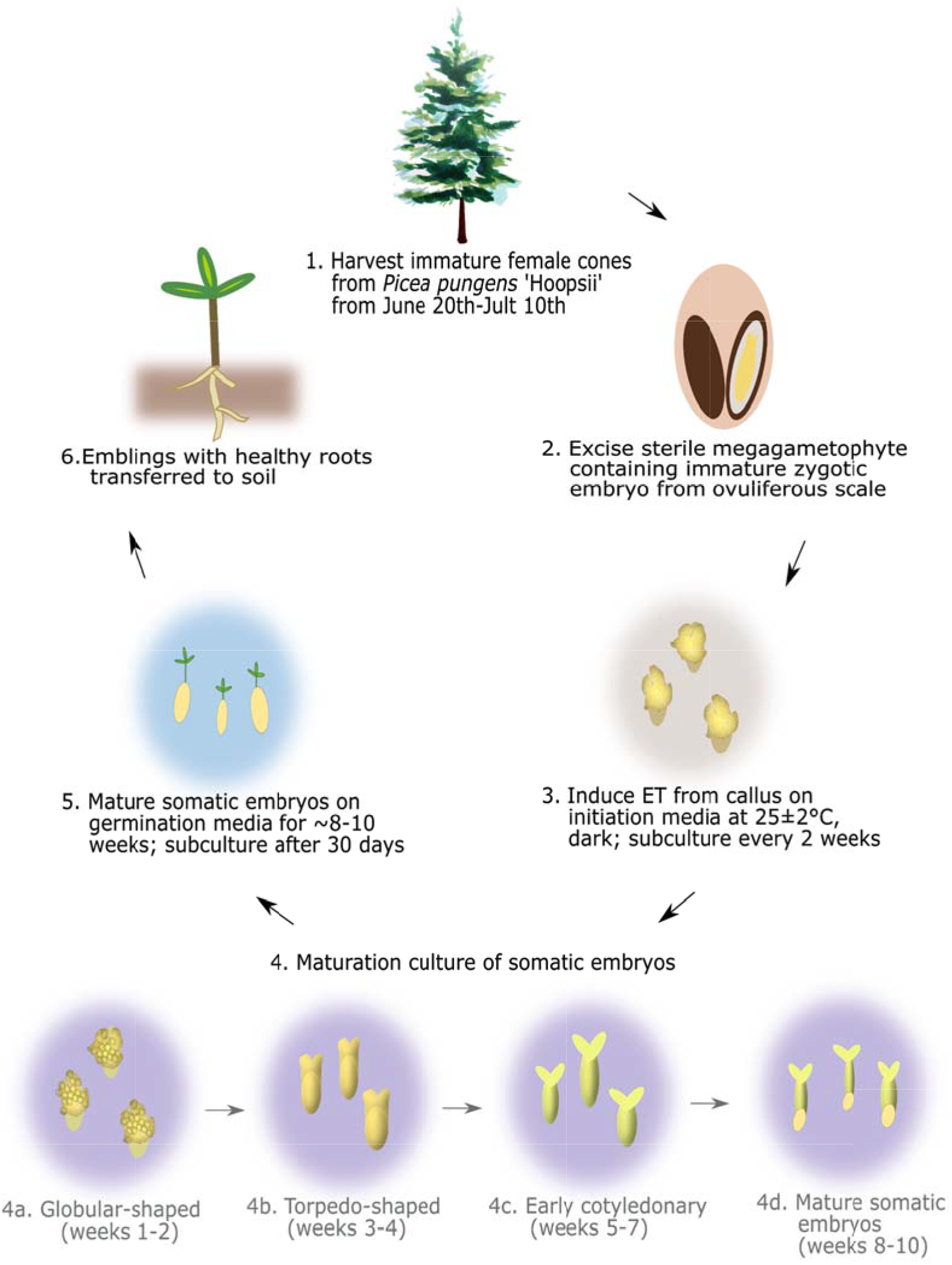
Overview of somatic embryogenesis using immature zygotic embryos from *Picea pungens* Hoopsii. *ET* embryonic tissue. Adapted from Scholthof et al (2018).

**Fig. 3.**
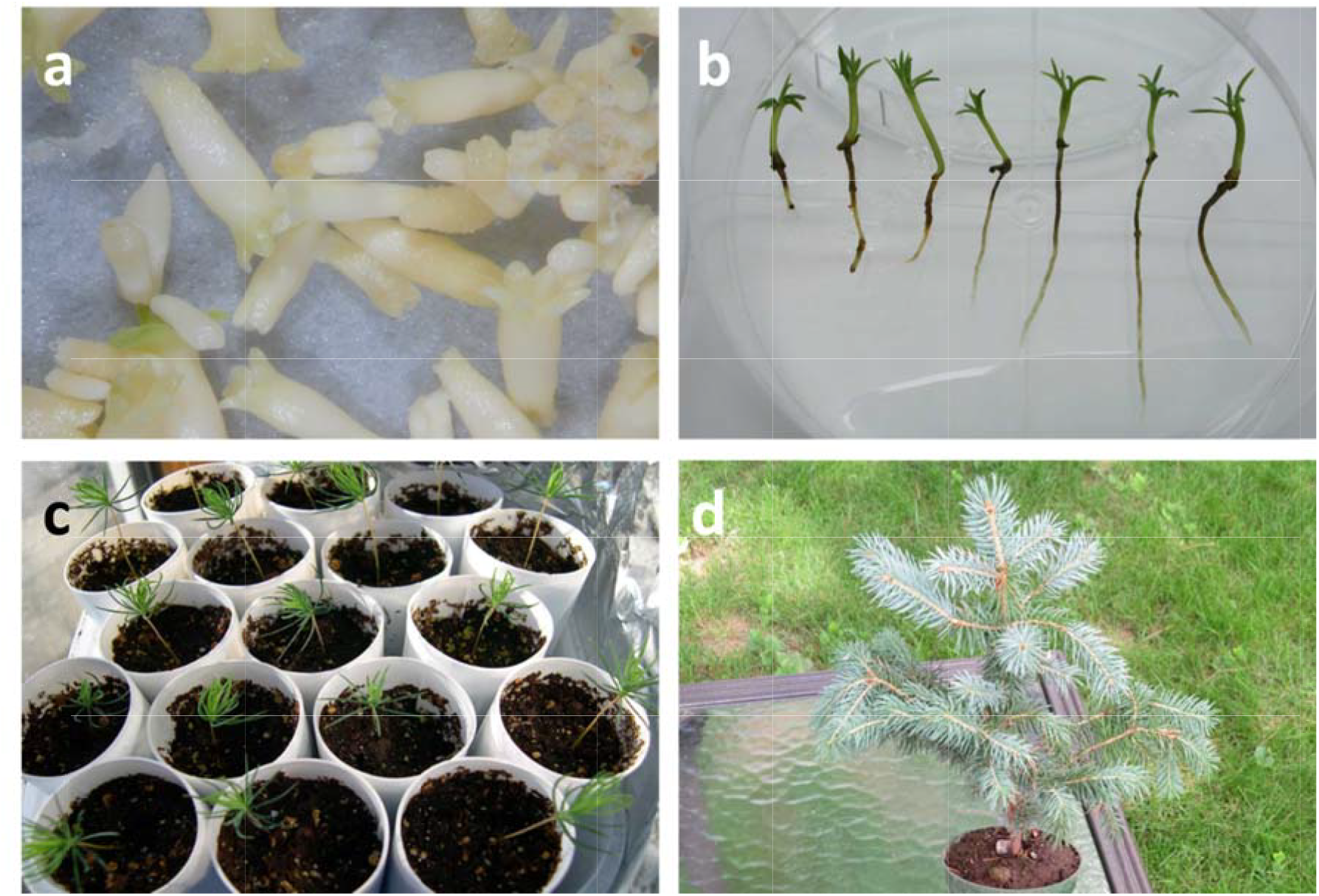
Development of *Picea pungens* Hoopsii somatic emblings **a** Mature somatic embryos **b** Eight-week-old somatic emblings on maturation media Hoopsii tree **c** Somatic emblings growing in soil **d** Three-month-old *P. pungens Hoopsii tree*

### Germination of somatic embryos

Cotyledons of somatic embryos produced chlorophyll and began to elongate within two days of being placed onto germination medium and exposed to light. The root cap turned pink and lateral roots began to develop. Once roots developed fully, the somatic emblings grew rapidly. By six weeks the shoots were 1.0–1.5 cm in height and roots were 0.5–2.0 cm long (Fig. 3b). At this stage the emblings were ready for transplantation to soil in the greenhouse. The conversion process (from immature zygotic embryo to a healthy rooted embling) was completed within eight to ten weeks. Overall, the average conversion rate from somatic embryos to emblings was 92.8%.

## Discussion

The difficulty of facilitating *Picea pungens* cultivar Hoopsii mass propagation through traditional methods has hindered development of this cultivar as a valuable product in the ornamental tree industry. Plant production via *in vitro* somatic embryogenesis is a potential answer to this problem. In this study, we spent approximately 20 months collecting and analyzing zygotic and somatic tissue samples from Hoopsii trees. The objective, which has been successfully fulfilled, was to establish a rapid and effective method of propagating this species via somatic embryogenesis, as summarized in Fig. 2.

ET initiation presented itself as the major challenge in establishing a protocol. The model conifer Norway spruce (*Picea abies*) successfully produces ET from mature somatic embryos (Bojarczuk et al. 2007), but we found that the optimal explant source was the precotyledonary immature zygotic embryo, collected from ~June 20 to ~July 10 (a develop-mental window of approximately two weeks) in eastern Ontario. Work in black pine (*Pinus nigra*) has demonstrated that the use of immature zygotic embryos enclosed within the intact megagametophytes is effective in producing ET that differentiate into viable somatic embryos for propagation (Salaj et al. 2019). Hoopsii megagametophytes containing small embryos collected prior to this window did not produce ET, nor did ovuliferous scales containing mature embryos collected after this window.

The average ET initiation frequency of zygotic embryos collected at this stage was 7.67%, which was the highest rate we attained from any explant source. The combination of 2,4-D and BAP in particular proved effective in producing ET callus from Hoopsii embryos, as it has for other conifer species (Pullman et al. 2016; Nunes et al. 2018). The combination of 2,4-D and BAP, however, has been used to successfully induce ET from Serbian spruce (*Picea omorika*), black spruce (*Picea mariana*), white spruce (*P. glauca*), Brewer spruce (*Picea breweriana*), and Norway spruce (*P. abies*), so the success of using these two plant growth regulators in conjunction may be general to *Picea*.

Following the initial establishment of a Hoopsii ET culture system and successful somatic embryo maturation, we found that SE-derived tissues could provide a virtually unlimited source of explants, providing the foundation for a solid Hoopsii propagation system. ET development under this system is also rapid; callus is produced approximately four weeks after plating immature zygotic embryos onto the initiation medium. This callus can be maintained for an additional four weeks, provided it is subcultured every two weeks onto fresh initiation medium. Conceivably, Hoopsii ET could also be cryostored using protocols optimized for spruces over the past several decades (Adams et al. 1994; Tikkinen et al. 2018a) and protect against shortages of embling-producing material if Hoopsii mass-propagation becomes a commercially viable endeavor in the future.

Maturation of ET into mature somatic embryos is a relatively robust and reliable step in this procedure (Fig. 3c, d). This phase required eight to ten weeks to produce the desired result of abundant Hoopsii trees (with a conversion rate of 92.8% (mature embryo to surviving embling)). Such a high yield exceeds yields resulting from culturing other species of conifer zygotic embryo for propagation: the conversion rate for an elite cultivar of Korean spruce (*Picea koraiensis* Nakai) has reached as high as ~25% using immature zygotic embryos as an explant source (Li et al. 2008). The black spruce (*P. mariana*) conversion rate has been improved to the point where it is close to reaching 50% survival. Both mature and immature zygotic embryos were used for *P. mariana* ET induction; while immature embryos exhibited higher ET rates, the study did not attach any importance to whether the emblings were derived from mature or immature embryos (Adams et al. 1994); the white spruce (*P. glauca*) conversion rate can approach 58% (although again no distinction was made between mature and immature zygotic embryos) (Stasolla and Yeung 1999); for Norway spruce, conversion rates range from as high as 77% when using immature zygotic embryos as an explant source (Tikkinen et al. 2018a), to as low as 3% when mature zygotic embryos were used as an explant source (Hazubska-Przybyl and Wawrzyniak 2017). The closest studied Hoopsii relative, blue spruce (*P. pungens* Engelmann), exhibited conversion rates of up to ~10%, although this particular protocol used zygotic embryos excised from mature seeds (Afele et al. 1992) rather than precotyledonary immature zygotic embryos. Mass propagation via somatic embryogenesis in Serbian spruce (*P. omorika*), Norway spruce (*P. abies*), Brewer spruce (*P. breweriana*) and white spruce (*P. glauca*) obtained conversion rates as high as 30% (Klimaszewska et al. 2001; Hazubska-Przybył 2008) – but again, only mature zygotic embryos were utilized. Our Hoopsii method is unique among *Picea* somatic embryogenesis protocols (with the exception of a recently developed *P. abies* protocol (Tikkinen et al. 2018a)) in that we harvest immature zygotic embryos that have not yet developed cotyledons, but are large enough to be excised from the megagametophyte. The use of immature zygotic embryos for the mass propagation of *P. pungens* Hoopsii and other spruces appears to be one crucial variable in obtaining conversion rates of over 30%. Recent work on *P. abies* has demonstrated that reducing nitrogen content in the germination medium can increase conversion rate and shorten the transfer time of the embling from *in vitro* growth conditions to the greenhouse (Tikkinen et al. 2018b), suggesting that the Hoopsii conversion rate can be further improved by following new techniques optimized in *P. abies*.

Moving forward, there are a myriad of steps that can be taken to ensure the future of the Hoopsii genotype. Mass propagation of desirable conifers could soon be performed at an industrial scale via the use of large-scale immersion bioreactors (Gonzalez-Cabrero et al. 2018). Interestingly, propagating Norway spruce via emblings increases the tree’s resistance to herbivory by the pine weevil (*Hylobius abietis* L.), a common conifer pest (Puentes et al. 2018). By exposing the embryos and young emblings to stress-inducing conditions during *in vitro* culturing, the emblings are plausibly ‘primed’ against herbivory. These benefits will become invaluable in the face of climate change, as some are predicting that *H. abietis* herbivory of conifers will increase as the global temperature increases (Haynes et al. 2014; Wainhouse et al. 2014). The establishment of a reliable tissue culture system also positions Hoopsii as a target for transgenesis; Norway spruce has been successfully transformed via *Agrobacterium tumefaciens*-mediated infection (Wenck et al. 1999), and in this vein Hoopsii and other conifers may become hosts for manipulation to improve insect resistance, or to possibly serve as mediators of atmospheric phytoremediation (Demone et al. 2018).

## Acknowledgments

The authors extend their gratitude for the funding provided by the Reductase Consortium, The Rockefeller Foundation, and NSERC (Natural Sciences and Engineering Research Council).

